# Allogeneic CRISPR-Engineered CAR-T Cells Drive Potent Antitumor Activity in Solid Tumors

**DOI:** 10.64898/2026.04.25.720815

**Authors:** Mingyu Huo, Dan Li, Nan Li, Alex Quan, Tianyuzhou Liang, Dan Henderson, Jason Sagert, Minh Pham, Luke Hanley, Kelly Maeng, Mikhala Eule, Mitchell Ho

## Abstract

Chimeric antigen receptor (CAR) T-cell therapy has shown limited efficacy in solid tumors, in part due to variability in autologous T cells derived from heavily pretreated patients with advanced disease. To address these constraints, we developed an off-the-shelf allogeneic CAR-T platform using CRISPR-Cas9-mediated genome editing in T cells from healthy donors to enable targeted CAR insertion at the *TRAC* locus with concurrent disruption of *B2M*. Using adeno-associated virus (AAV) delivery, we designed CAR-T cells targeting glypican-2 (GPC2) and glypican-3 (GPC3), emerging antigens expressed in pediatric and adult solid tumors. Genome-edited allogeneic CAR-T cells exhibited potent, antigen-specific cytotoxicity across multiple tumor models. GPC2-directed allogeneic CAR-T cells demonstrated enhanced or comparable activity relative to conventional lentiviral CAR-T cells in neuroblastoma models and mediated tumor regression with prolonged survival in preclinical models. Notably, repeated dosing augmented antitumor efficacy without evidence of toxicity, supporting multi-dose regimens for solid tumors. Similarly, GPC3-targeted allogeneic CAR-T cells based on a single-domain antibody showed robust activity against hepatocellular carcinoma cells in vitro and in vivo. These findings establish a scalable, genome-engineered allogeneic CAR-T strategy with strong therapeutic potential and support the clinical development of off-the-shelf cell therapies for pediatric and adult solid tumors.

## Introduction

Chimeric antigen receptor (CAR) T-cell therapy has transformed the treatment landscape for hematologic malignancies.^1^ However, the clinical success of conventional CAR-T therapies has not translated effectively to solid tumors and remains constrained by reliance on autologous T cells derived from heavily pretreated patients with advanced disease. These cells often exhibit impaired fitness, limited expansion capacity, and functional heterogeneity, contributing to variable therapeutic efficacy. In addition, individualized manufacturing is time-intensive, costly, susceptible to failure, particularly in patients with lymphopenia or compromised T-cell quality following prior radiotherapy and chemotherapy.^2-5^ Collectively, these limitations highlight the need for alternative strategies to enable scalable, consistent, and effective CAR-T therapies.

To address these challenges, allogeneic CAR-T cell therapies derived from healthy donors or engineered cell sources have emerged as a promising alternative. In contrast to autologous approaches, these “off-the-shelf” products can be manufactured at scale, cryopreserved, and delivered on demand to patients, enabling greater consistency and reduced production timelines and costs. Moreover, the use of healthy donor T cells may enhance cellular fitness and reduce exhaustion thereby improving therapeutic potency.^6,7^

Early-phase clinical trials of allogeneic CAR-T cells targeting CD19,^8,9^ CD70,^10^ and B-cell maturation antigen (BCMA)^11^ have demonstrated encouraging response rates, without reports of graft-versus-host disease (GvHD). Despite these advances, GvHD and host-versus-graft disease (HvGD) remain significant barriers to broader clinical application. To mitigate GvHD, genome editing tools such as CRISPR-Cas9 have been used to disrupt the endogenous T cell receptor (TCR), most commonly by targeting the T cell receptor alpha constant (*TRAC*) locus.^12,13^ Strategies to prevent HvGD include lymphodepletion, human leukocyte antigen (HLA) editing, and immune evasion techniques.^13-15^

Glypicans (GPCs) are a family of heparan sulfate proteoglycans anchored to the cell surface via glycosylphosphatidylinositol anchorage.^16-18^ Among them, GPC2 and GPC3 are oncofetal antigens highly expressed in neuroblastoma and hepatocellular carcinoma, respectively, making them emerging targets for immunotherapy.^17,19-21^ Our laboratory has previously developed high-affinity antibodies against these antigens, including the GPC2-specific CT3 and the GPC3-specific hYP7 and single-domain antibody HN3, which demonstrate robust antitumor activity when incorporated into CAR-T cell formats.^22,23^

In this study, we developed a genome-engineered allogeneic CAR-T platform using CRISPR-Cas9 to disrupt *TRAC* and beta-2-microglobulin (*B2M*), thereby eliminating endogenous TCR expression and reducing MHC I-mediated immune recognition. Concurrently, we used adeno-associated virus (AAV) -mediated homology-directed repair to enable CAR insertion at the *TRAC* locus. Using this strategy, we engineered CAR-T cells targeting GPC2 and GPC3 and systematically optimized CAR design, including antigen-binding single chain variable fragment (scFv) orientation and co-stimulatory domains. Selected CAR-T cells, CT3vHvL-CD28 (1528) for GPC2 and HN3-CD28 (1524) for GPC3, demonstrated potent and antigen-specific antitumor antitumor activity across multiple preclinical murine models. Notably, repeated dosing of allogeneic CAR-T cells showed no evidence of toxicity. These findings support the feasibility of a scalable, off-the-shelf CAR-T strategy for the treatment of solid tumors.

## Results

### Generation and characterization of GPC2-targeted allogeneic CAR-T cells

Electroporation was used to deliver Cas9 protein and single-guide RNAs (sgRNAs) targeting the *TRAC* and *B2M* loci, followed by transduction with a donor template DNA to insert the corresponding CAR construct into the *TRAC* locus (Fig. 1A). We designed four CAR constructs containing the CT3 antibody in two scFv orientations (vH-vL and vL-vH), each paired with either CD28 or 4-1BB co-stimulatory domains, specifically: CT3vHvL-CD28 (1528), CT3vHvL-4-1BB (1528b), CT3vLvH-CD28 (1529), CT3vLvH-4-1BB (1529b) (Fig. 1B). Flow cytometry was used to detect CAR expression (knock-in efficiency), which ranged from 43.8% to 74%, with 1528b achieving the highest efficiency (74%) and 1529 the lowest (43.8%) (Fig. 1C). *TRAC* and *B2M* knockout efficiencies were consistently high across all constructs, 98-99% for *TRAC* and 75-80% for *B2M*, with no significant differences among the four groups (Fig. 1D).

**Figure 1.**
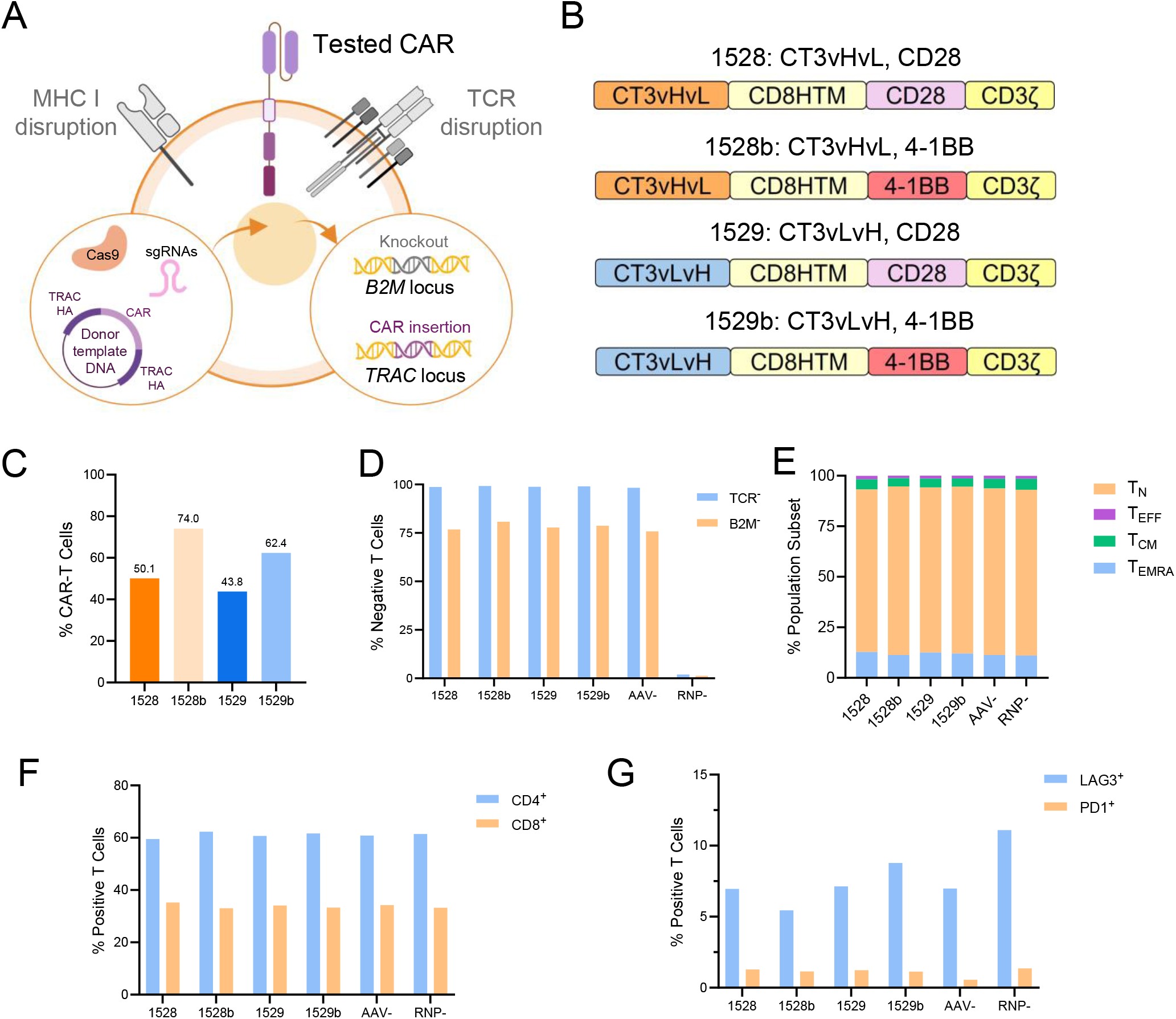
Generation and characterization of allogeneic CAR-T cells targeting GPC2. (A) Schematic workflow for generating CRISPR-engineered allogeneic CAR-T cells via CAR knock-in at the *TRAC* locus and *B2M* knock-out. AAV vectors were used as donor templates to deliver the CAR cassette. TRAC HA, TRAC homology arm. (B) Schematics of the four CAR constructs evaluated in this study. (C) Knock-in efficiency (percentage of CAR^+^ cells) of the four allogeneic CAR-T cells. (D) Knock-out efficiency of *TRAC* and *B2M* assessed by flow cytometry as the percentage of marker-negative cells. AAV-(Cas9 and sgRNA without donor) and RNP-(CRISPR Ribonucleoprotein-, electroporation buffer only) were used as negative controls. (E) Differentiation phenotypes of allogeneic CAR-T cells, including naïve (T_N_), effector (T_EFF_), central memory (T_CM_), and effector memory CD45RA^+^ (T_EMRA_) subsets. (F) Proportions of CD4^+^ and CD8^+^ subsets among allogeneic CAR-T cells. (G) Expression levels of exhaustion markers LAG3 and PD-1 on allogeneic CAR-T cells.

We next assessed T cell differentiation using flow cytometry. All allogeneic CAR-T cells and control groups exhibited similar phenotypic distributions, with approximately 82% naïve T cells (T_N_), 12% terminally differentiated effector memory T cells (T_EMRA_), 4.5% central memory T cells (T_CM_), and 1.5% effector T cells (T_EFF_) (Fig. 1E). CD4/CD8 composition was also consistent across groups, with around 60% CD4^+^ and 33% CD8^+^ CAR-T cells (Fig. 1F). To evaluate the exhaustion profile, we measured lymphocyte-activation gene 3 (LAG3) and programmed cell death protein 1 (PD-1) expression. All four allogeneic CAR-T cell types exhibited low and comparable levels of exhaustion markers, with 5-8% LAG3^+^ and around 1% PD-1^+^ cells (Fig. 1G).

### GPC2-targeted allogeneic CAR-T cells exhibit antigen-specific cytotoxicity and cytokine production in neuroblastoma models

To evaluate the efficacy of allogeneic CAR-T cells, we conducted the *in vitro* cytotoxicity assay using the GPC2-high neuroblastoma cell line IMR-32 and the GPC2-negative renal carcinoma cell line A498 as a control. All four anti-GPC2 allogeneic CAR-T cells demonstrated comparable cytotoxicity against IMR-32 tumor cells (Fig. 2A). In contrast, none of the constructs exhibited killing activity against the GPC2-negative A498 cells, indicating a lack of off-target or non-specific cytotoxicity (Fig. 2B).

**Figure 2.**
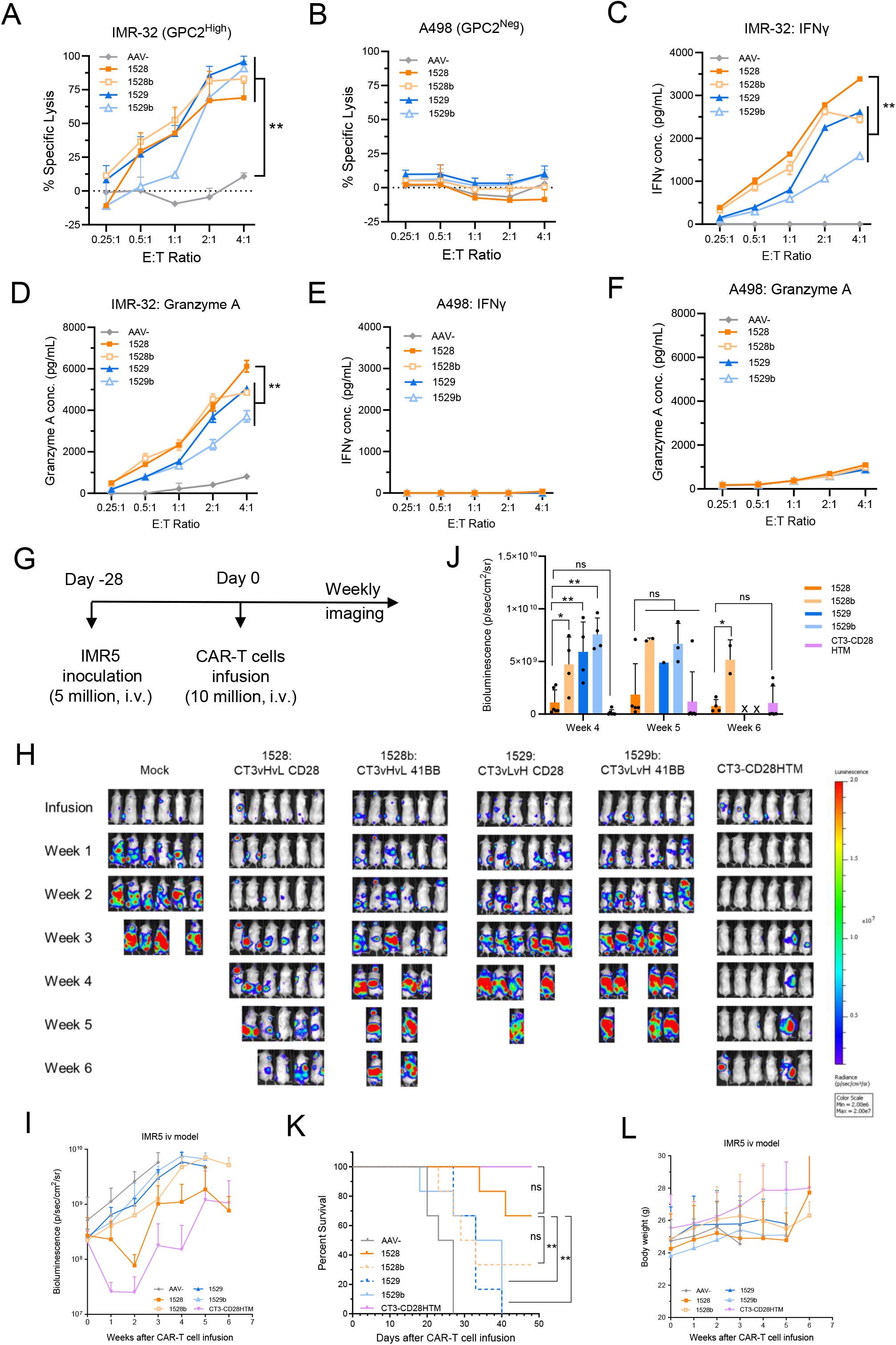
*In vitro* functionality and *in vivo* efficacy of anti-GPC2 allogeneic CAR-T cells against neuroblastoma tumors. (A-B) Cytotoxicity assays showing tumor cell killing by allogeneic CAR-T cells after 24-hour co-culture with IMR-32 (A) and A498 (B) cells. (C-D) IFN-γ (C) and Granzyme A (D) secretion by allogeneic CAR-T cells in response to IMR-32 co-culture. (E-F) IFN-γ (E) and Granzyme A (F) secretion by allogeneic CAR-T cells in response to A498. All *in vitro* experiments were conducted in triplicate. (G) Schematic of the *in vivo* IMR5 xenograft model: NSG mice were inoculated with 5 × 10□ IMR5 tumor cells for 28 days, followed by intravenous infusion of 1 × 10□ CAR-T cells. N = 6 mice per group. (H) Representative tumor bioluminescence images. (I) Tumor growth curves over time. (J) Quantification of tumor bioluminescence signals. (K) Kaplan–Meier survival analysis. (L) Mouse body weight monitoring. Data are represented as mean ± SD. ^ns^P>0.05, *P<0.05, **P<0.01; ns, not significant.

We also assessed cytokine secretion by the allogeneic CAR-T cells upon co-culture. CT3vHvL-CD28 (1528) CAR-T cells secreted significantly higher levels of interferon-γ (IFN-γ) and Granzyme A in response to IMR-32 stimulation among the four allogeneic CAR-T cell groups (Fig. 2C, 2D). No significant cytokine release was observed when CAR-T cells were co-cultured with the A498 cells, confirming antigen-dependent activation (Fig. 2E, 2F). Together, these data demonstrate that all four anti-GPC2 allogeneic CAR-T cells exhibit similar antigen-specific cytotoxicity in cell culture, with higher cytokine production in the CT3vHvL-CD28 (1528) group.

### CT3vHvL-CD28 outperforms in the screening of anti-GPC2 allogeneic CAR-T cells in a metastatic neuroblastoma model

To assess allogeneic CAR-T cells in preclinical animal models, we tested the four anti-GPC2 allogeneic CAR-T cells in a neuroblastoma metastasis mouse model. Mice were intravenously injected with IMR5 neuroblastoma tumor cells and treated with CAR-T or mock T cells when tumor signals reached approximately 5 × 10□ p/sec/cm^2^/sr (Fig. 2G). CT3 CAR-T cells generated by lentiviral transduction served as a positive control.

Although none of the allogeneic CAR-T cells matched the efficacy of the lentiviral CT3 CAR-T cells, CT3vHvL-CD28 (1528) showed better performance among the four allogeneic CAR-T cell groups. Mice in the CT3vHvL-CD28 (1528) group had slower tumor progression (Fig. 2H-J), better survival (Fig. 2K), and maintained higher body weight (Fig. 2L) compared to those receiving CT3vHvL-4-1BB (1528b), CT3vLvH-CD28 (1529), or CT3vLvH-4-1BB (1529b). By the study endpoint, four of six mice treated with CT3vHvL-CD28 (1528) CAR-T cells remained alive, whereas only two or none survived in the other allogeneic CAR-T cell groups (Fig. 2K).

### Allogeneic CT3vHvL-CD28 CAR-T cells demonstrate broad and antigen-specific activity across neuroblastoma cell lines

We further validated the performance of CT3vHvL-CD28 (1528) CAR-T cells *in vitro* using five additional neuroblastoma cell lines: GPC2-high (G10, NBEB, IMR5) and GPC2-low (SKNSH, LAN1), as well as a GPC2-knockout line (IMR5-KO). Both CT3vHvL-CD28 (1528) and lentiviral CT3 CAR-T cells exhibited strong cytotoxicity against SKNSH and NBEB (Fig. 3A-B). Notably, CT3vHvL-CD28 (1528) CAR-T cells showed better killing of G10, IMR5, and LAN1 than lentiviral CT3 CAR-T cells (Fig. 3C-E). No cytotoxicity was observed against IMR5-KO cells for either group, confirming antigen specificity (Fig. 3F). These results demonstrate that CT3vHvL-CD28 (1528) allogeneic CAR-T cells have similar or slightly better efficacy than lentiviral CT3 CAR-T cells *in vitro*.

**Figure 3.**
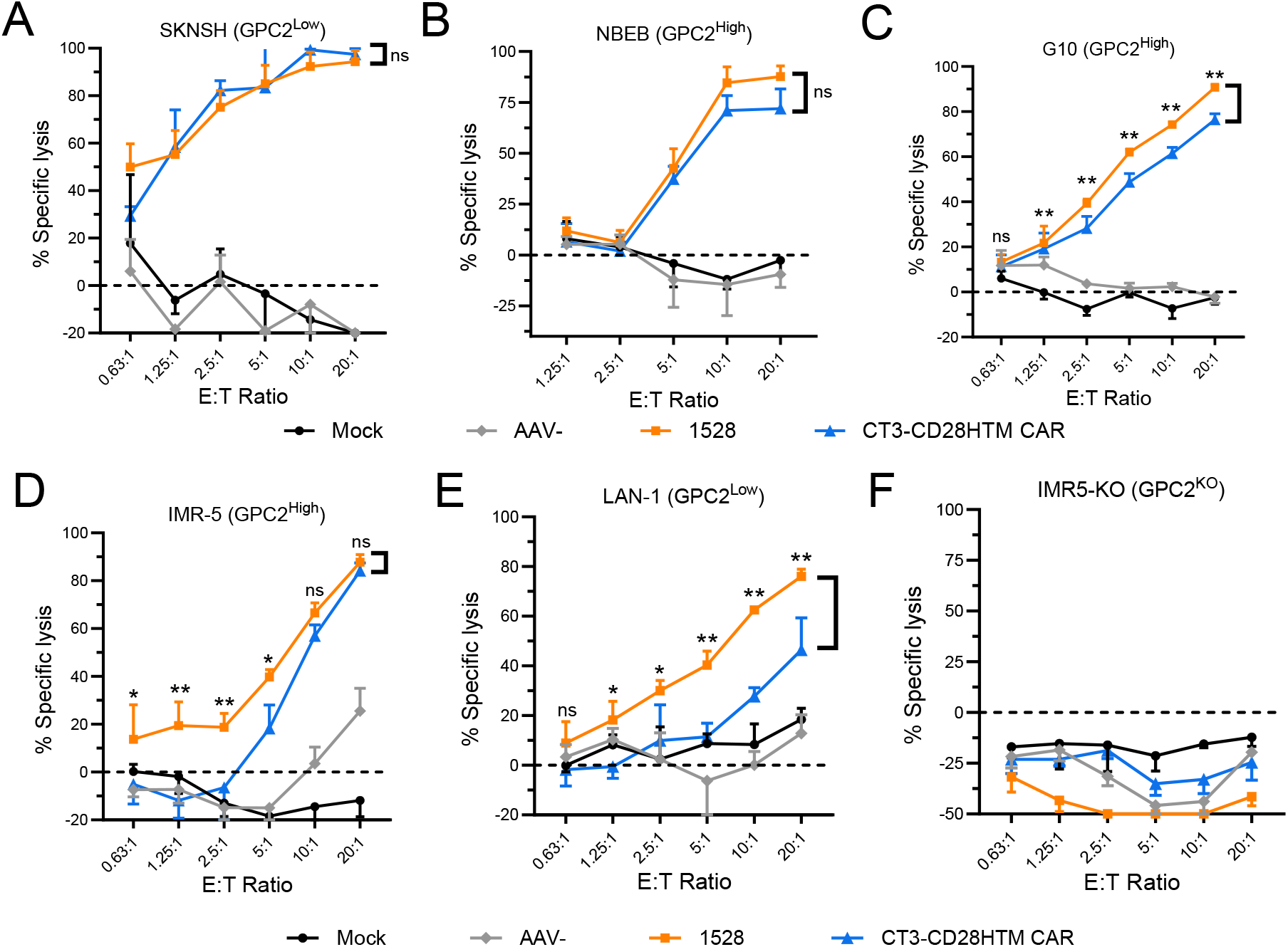
*In vitro* cytotoxicity of allogeneic CT3vHvL-CD28 CAR-T cells against a panel of neuroblastoma cell lines. (A-F) Cytotoxicity assays of allogeneic CT3vHvL-CD28 (1528) and lentiviral CT3-CD28HTM CAR-T cells against SKNSH (A), NBEB (B), G10 (C), IMR5 (D), LAN1 (E), and GPC2-knockout IMR5 cells (F). AAV- and Mock groups were used as controls. Tumor killing was assessed at various effector-to-target (E:T) ratios. Data represent three independent experiments and are represented as mean ± SD. ^ns^P>0.05, *P<0.05, **P<0.01.

### Allogeneic CT3vHvL-CD28 CAR-T cells demonstrate a favorable safety profile in a high-tumor burden model

To test the performance of CT3vHvL-CD28 (1528) CAR-T cells against larger tumors, we injected mice with IMR5 cells intravenously and administered 10 million CAR-T cells when tumor signals reached around 1 × 10□ p/sec/cm^2^/sr (Fig. 4A). CT3vHvL-CD28 (1528) CAR-T cells induced tumor regression and slower growth compared to controls. While these differences were not statistically significant, four out of seven mice consistently showed reduced tumor burden until week three (Fig. 4B-D). At the end of the experiment, two of seven mice in the CT3vHvL-CD28 (1528) CAR-T cell group remained alive, compared to one in the control group (Fig. 4E). The surviving CT3vHvL-CD28 CAR-T cell-treated mice also maintained higher body weights, suggesting improved physical condition (Fig. 4F).

**Figure 4.**
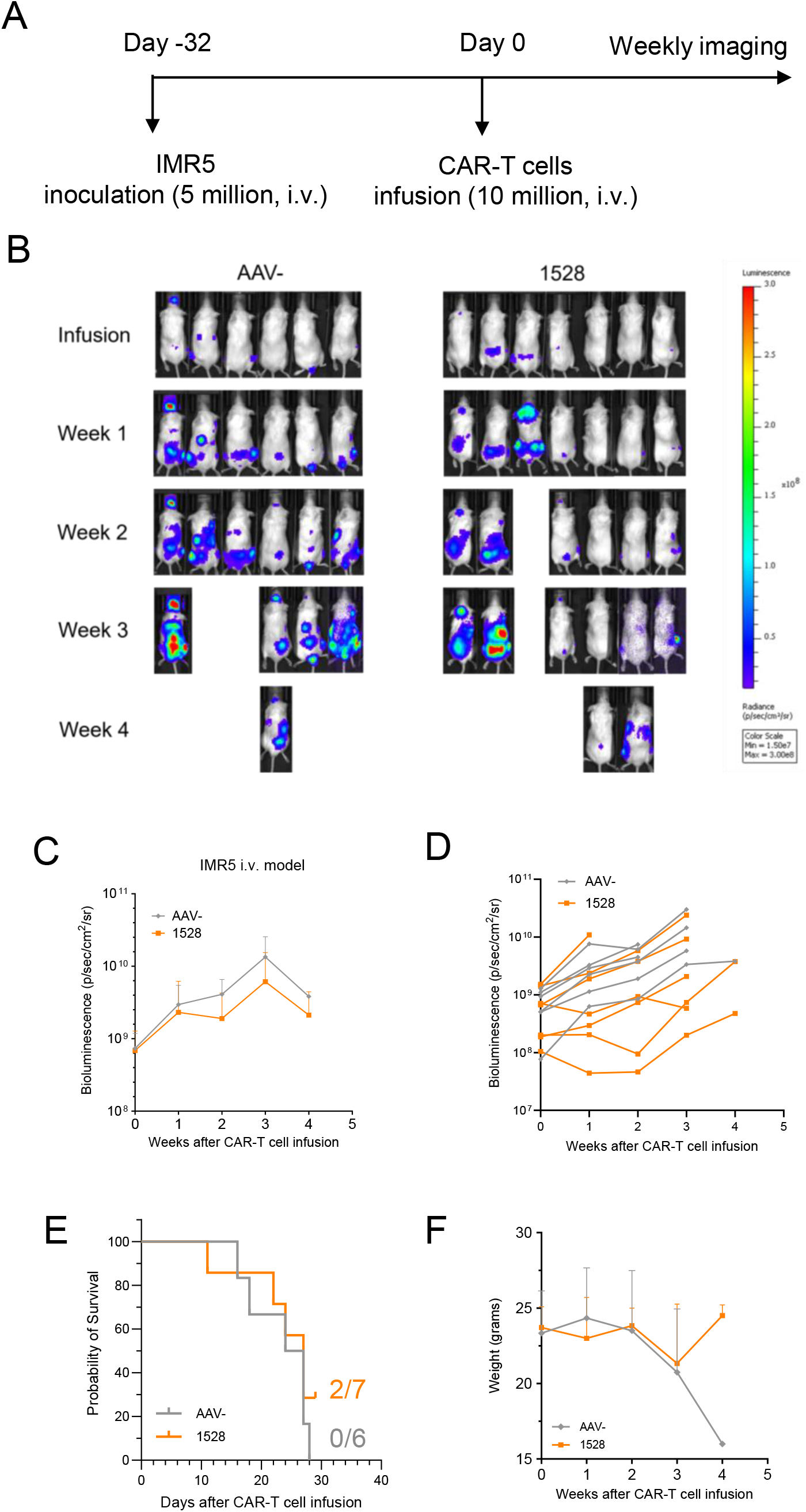
Allogeneic CT3vHvL-CD28 CAR-T cells are effective in an established mouse model of neuroblastoma. (A) Schematic of the *in vivo* experiment: NSG mice were inoculated with 5 × 10□ IMR5 cells for 32 days, then received 1 × 10□ AAV- or allogeneic CT3vHvL-CD28 (1528) CAR-T cells via tail vein injection. (B) Representative tumor bioluminescence images. (C) Average tumor growth curves. (D) Individual tumor growth trajectories. (E) Kaplan-Meier survival curves. (F) Mouse body weight monitoring throughout the study. N = 6 or 7 mice per group. Data are represented as mean ± SD.

### Repeated dosing of allogeneic CAR-T cells enhances antitumor activity without added toxicity

To evaluate the feasibility of multiple treatments, we used the same neuroblastoma metastasis model with smaller tumor burdens. Mice received three doses of either allogeneic CT3vHvL-CD28 (1528) CAR-T cells or control T cells on days 0, 16, and 30 (Fig. 5A). CT3vHvL-CD28 (1528) CAR-T cells treatment led to significant tumor regression by weeks three and four compared to controls (Fig. 5B-D). By day 50, four of seven mice in the CT3vHvL-CD28 (1528) CAR-T cells group were still alive, while all mice in the control group had died by day 34 (Fig. 5E). The surviving mice maintained or gained weight and appeared in good physical condition (Fig. 5F). These results indicate that CT3vHvL-CD28 (1528) CAR-T cells are safe and effective when administered in multiple doses and are particularly efficacious against smaller tumors.

**Figure 5.**
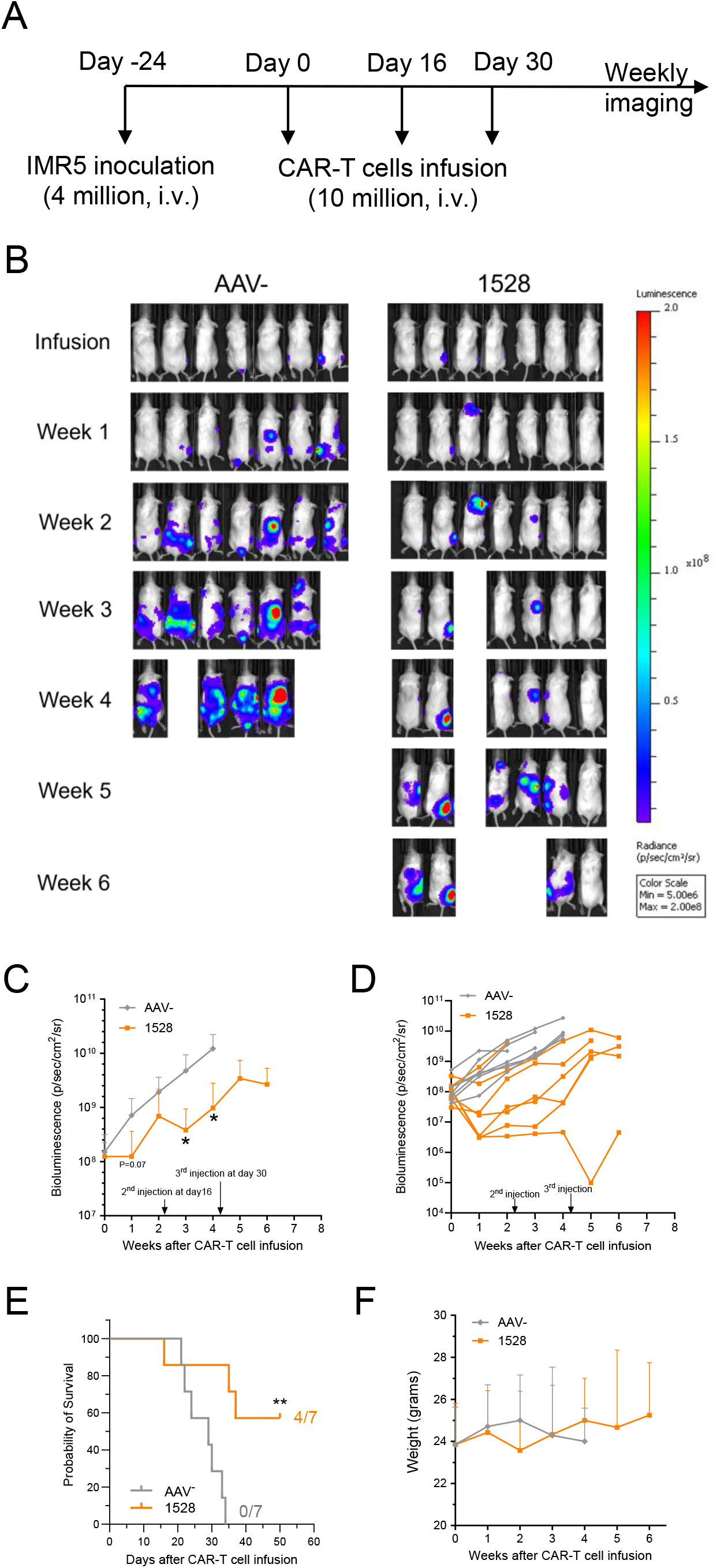
Multiple injections of allogeneic CT3vHvL-CD28 CAR-T cells enhance efficacy in early-stage neuroblastoma tumors. (A) Experimental design: NSG mice were injected with 4 × 10□ IMR5 tumor cells for 24 days, followed by three intravenous doses of 1 × 10□ AAV- or allogeneic CT3vHvL-CD28 (1528) CAR-T cells. (B) Representative tumor bioluminescence images. (C) Average tumor growth curves. (D) Individual tumor growth trajectories. (E) Kaplan-Meier survival curves. (F) Body weight monitoring of treated mice. N = 7 mice per group. Data are represented as mean ± SD. ^ns^P>0.05, *P<0.05, **P<0.01.

### Screening of GPC3-targeted allogeneic CAR-T cells identifies lead candidates for hepatocellular carcinoma

To apply this strategy to hepatocellular carcinoma, we designed eight anti-GPC3 allogeneic CAR-T cells, incorporating two antibody fragments (HN3 V_H_ and hYP7 scFv) in various scFv orientations and co-stimulatory domains (CD28 or 4-1BB). The constructs include HN3-CD28 (1524), HN3-4-1BB (1524b), hYP7-vHvL-CD28 (1525), hYP7-vHvL-4-1BB (1525b), hYP7-vLvH-CD28 (1526), hYP7-vLvH-4-1BB (1526b), IgG4CH2CH3-CD28 (1531), and IgG4CH2CH3-4-1BB (1531b) (Fig. 6A).

**Figure 6.**
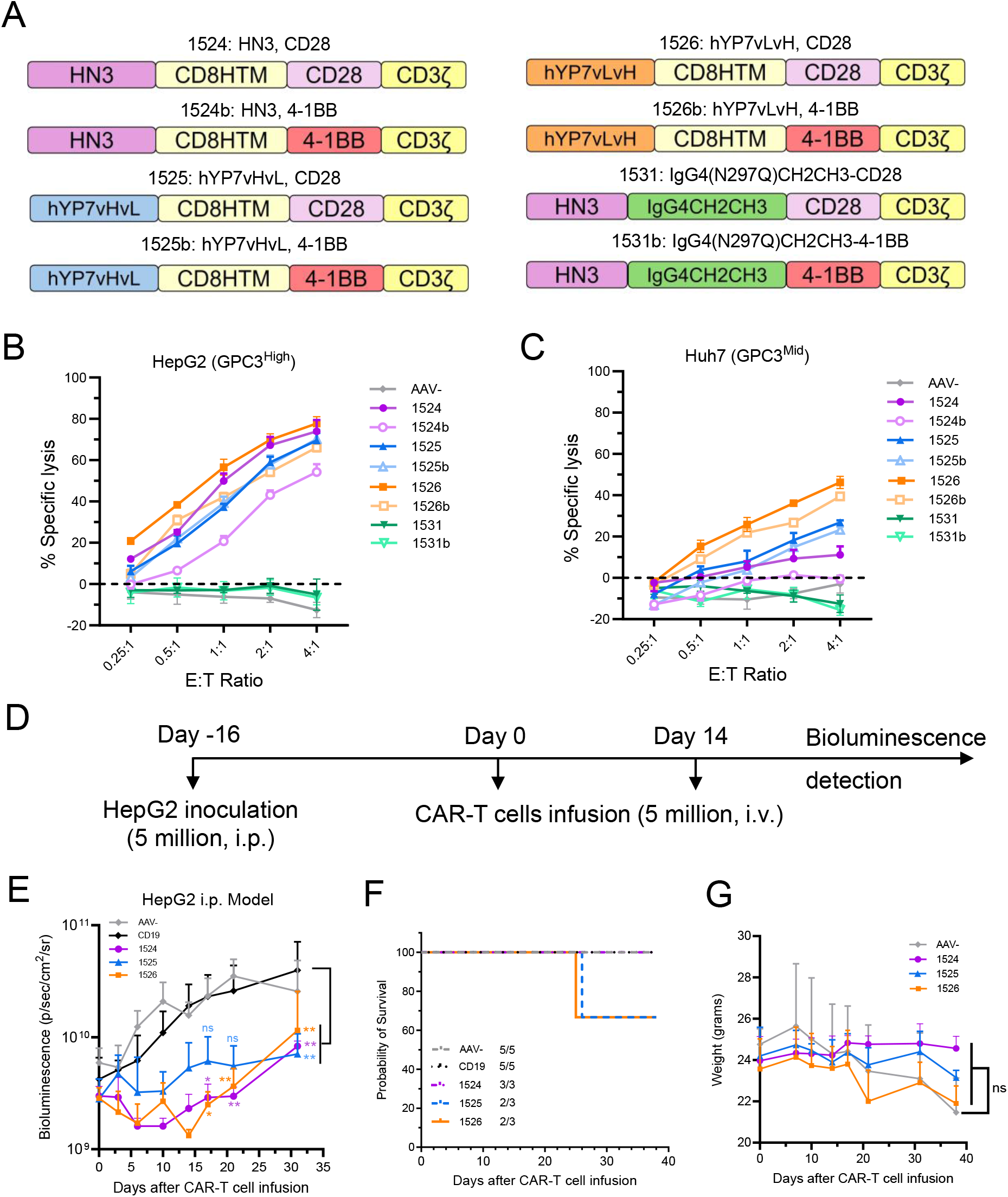
Characterization of GPC3-targeting allogeneic CAR-T cells against hepatocellular carcinoma *in vitro* and *in vivo*. (A) Schematic representation of the eight CAR constructs targeting GPC3. (B-C) *In vitro* cytotoxicity assays of the eight GPC3-targeting allogeneic CAR-T cells against hepatocellular carcinoma cell lines, HepG2 (B) and Huh7 (C) cells, at multiple E:T ratios. AAV-was used as the control. N = 3 independent experiments. (D) *In vivo* experimental schematic: NSG mice were inoculated with 5 × 10□ HepG2 cells, followed by two intravenous doses of 5 × 10□ AAV- or CAR-T cells at Day 0 and Day 14 post-tumor engraftment. (E) Tumor growth curves. (F) Kaplan-Meier survival analysis. (G) Mouse body weight over the course of treatment. N = 5 mice per group. Data are represented as mean ± SD. ^ns^P>0.05, *P<0.05, **P<0.01.

We performed *in vitro* killing assays using two GPC3-positive hepatocellular carcinoma cell lines, HepG2 and Huh7. Among the allogeneic CAR-T cells, HN3-CD28 (1524), hYP7vHvL-CD28 (1525), and hYP7vLvH-CD28 (1526) CAR-T cells demonstrated superior cytotoxic activity compared to the other groups (Fig. 6B-C). Next, we established a HepG2 xenograft mouse model to evaluate the efficacy of these three allogeneic CAR-T cells (Fig. 6D). All three allogeneic CAR-T cell treatments led to significant tumor regression compared to CD19 CAR-T controls. Although the differences between the three allogeneic CAR groups were not statistically significant, HN3-CD28 (1524) CAR-T cells consistently outperformed the others, resulting in smaller tumor volumes (Fig. 6E), extended survival (Fig. 6F), and higher body weight maintenance in treated mice (Fig. 6G). These results indicate that HN3-CD28 (1524), hYP7vHvL-CD28 (1525), and hYP7vLvH-CD28 (1526) CAR-T cells are the most promising GPC3-targeted CAR designs, with HN3-CD28 (1524) emerging as the lead candidate based on both *in vitro* and *in vivo* efficacy.

## Discussion

In this study, we establish a CRISPR-engineered, AAV-enabled allogeneic CAR-T platform for the development of scalable, “off-the-shelf” cell therapies targeting solid tumors. By site-specifically integrating GPC2- and GPC3-directed CAR constructs into the *TRAC* locus while simultaneously disrupting *B2M*, we achieved precise genomic engineering that supports consistent CAR expression and reduced alloreactivity. The resulting allogeneic CAR-T cells exhibited potent, antigen-specific cytotoxicity in vitro and robust antitumor activity in vivo, coupled with a favorable safety profile. Importantly, repeated administration of up to three doses was well tolerated, with treated animals maintaining good physical condition throughout, highlighting the feasibility of multi-dose regimens as a strategy to enhance therapeutic efficacy in solid tumors.

To evaluate safety under immunosuppressive conditions, we used a mouse model bearing a large tumor burden. Under high antigen load and an immunosuppressive microenvironment, allogeneic CAR-T treatment led to reduced tumor burden, prolonged survival, and improved overall physical condition compared to control groups. We further demonstrated that allogeneic CAR-T cells are effective in a setting of smaller tumor burdens, with repeated high-dose administrations leading to significant tumor regression and enhanced survival, supporting the feasibility of multi-dose regimens.

Our comparison of CAR constructs revealed that the CD28 co-stimulatory domain consistently outperformed its 4-1BB counterpart when paired with the same antibody and scFv orientation. Among the constructs tested, HN3-CD28 (1524), hYP7vLvH-CD28 (1526), and CT3vHvL-CD28 (1528) each showed superior performance within their respective antibody groups, HN3, hYP7, and CT3, demonstrating the importance of CAR design in optimizing therapeutic efficacy. Interestingly, these top-performing constructs differed in scFv orientation: hYP7vLvH-CD28 (1526) employed a VL-VH configuration, while CT3vHvL-CD28 (1528) used VH-VL. Because scFv orientation influences folding, antigen binding, stability, and expression,^24,25^ our findings underscore the critical role of scFv configuration in CAR-T design and support the need to optimize this variable in future CAR-T development.

Despite these promising results, allogeneic CT3vHvL-CD28 CAR-T cells did not perform as well as traditional lentivirally transduced CT3 CAR-T cells in neuroblastoma models. Given NSG mice lack functional natural killer (NK) cells, this difference is unlikely to reflect NK cell-mediated clearance resulting from *B2M* knockout. Instead, we speculate that disruption of the endogenous TCR may impair long-term persistence as TCR signaling has been implicated in promoting homeostatic survival and memory formation.^26,27^ These findings highlight a potential trade-off between minimizing alloreactivity and preserving long-term CAR-T cell fitness. Future Strategies to enhance persistence and function may include cytokine engineering (e.g., IL-15 or IL-7/21 co-expression) ^28-30^ and the incorporation of co-stimulatory ligands such as 4-1BBL.^31^ Additional genetic modifications, such as PD-1 knockout to mitigate exhaustion or HLA-E overexpression to reduce susceptibility to NK cell-mediated clearance, may further enhance *in vivo* performance.^32,33^

Taken together, our findings establish a CRISPR-engineered, AAV-enabled technology for generating functional, “off-the-shelf” allogeneic CAR-T cells targeting GPC2 and GPC3. This strategy directly addresses critical limitations of autologous CAR-T therapies and offers a promising alternative for patients with advanced disease or poor-quality T cells. Continued optimization of gene editing strategies and persistence-enhancing modifications will be essential for maximizing the clinical potential of next-generation allogeneic CAR-T therapies.

## Materials and methods

### Culture and generation of allogeneic CAR-T cells

CAR constructs were synthesized and cloned into an AAV6 transfer plasmid backbone. All constructs included a CD8 transmembrane domain in tandem with either a 4-1BB or CD28 costimulatory linked to a CD3ζ signaling domain. Gene editing and cell preparation were performed using standard techniques as described previously.^34^ Briefly, human peripheral blood mononuclear cells (Charles River Laboratories) were thawed and the T cells were activated with TransAct beads (Miltenyi Biotec) for three□days in T cell media (X-VIVO® 15 Serum-free Hematopoietic Cell Medium (Lonza) containing human serum (BioIVT), hIL-2 (Miltenyi Biotec) and hIL-7 (Sartorius)). After activation, the T cells were electroporated with Cas9 protein and single-guide RNAs (sgRNAs) targeting the *TRAC* and *B2M* loci and then transduced with a recombinant AAV6 vector containing donor template DNA for insertion of the corresponding CAR construct into the *TRAC* locus. Following electroporation and transduction, the CAR T cells were expanded for seven□days in the T cell media and cryopreserved.

### Flow cytometry

All flow cytometry experiments were conducted and analyzed using the SONY ID7000 (Sony Biotechnology) and the FlowJo software (BD Biosciences). The anti-EGFR human monoclonal antibody cetuximab (Erbitux) and a goat-anti-human IgG antibody conjugated with APC (Jackson ImmunoResearch) were used to stain CAR for analyzing knock-in efficiencies. The secondary antibody-only control served as the group for accounting for any nonspecific background signal. Anti-human TCRαβ (BD Biosciences) and anti-human HLA-ABC (BD Biosciences) were used to check knock-out efficiencies. Anti-human CD4 (BD Biosciences), anti-human CD8 (Biolegend), anti-human LAG3 (Biolegend), and anti-human PD1 (Invitrogen) were used to study CAR-T cells’ subpopulation and exhaustion markers. The CAR-T cell memory phenotype was analyzed using anti-human CCR7 (BD Biosciences), anti-human CD45RA (BD Biosciences), and anti-human CD45 (Biolegend). The CAR-T cells differentiation subsets consisted of naïve T cells (T_N_ cells; CD45RA+CCR7+), central memory T cells (T_CM_ cells; CD45RA-CCR7+), effector T cells (T_EFF_ cells; CD45RA-CCR7-), and terminally differentiated effector memory T cells (T_EMRA_ cells; CD45RA+CCR7-).

### *In vitro* cell killing assays and ELISA

Mock, AAV- and CAR-T cells were co-cultured with luciferase-expressing target cells at defined effector-to-target (E:T) ratios for 24 hours in triplicate wells. The tumor target cells used include GPC2-high-expression tumor cell lines, IMR-32, G10, NBEB, and IMR5; GPC2-low-expression tumor cell lines, SKNSH and LAN-1; GPC2-negative-expression cell lines, IMR5-KO and A498; GPC3-high-expression tumor cell line, HepG2; and GPC3-medium-expression tumor cell line, Huh7. G10 is a transfected A431 cell line stably expressing human GPC2. The supernatant was collected and centrifuged to remove cells and debris, and ELISA kits were used to detect IFNγ and Granzyme A following the manufacturer’s protocol (R&D). Tumor cells were lysed by the passive lysis buffer (E1941, Promega) at room temperature. Add luciferase assay reagent (E1501, Promega) to each sample and then measure the light produced using the SpectraMax (Molecular Devices) plate reader. Luminescence readings were normalized to target-only control wells (no T cells).

### Animal studies

The animal studies were under the protocol (LMB-059) approved by the Institutional Animal Care and Use Committee at the NIH. Five-week-old female NOD/SCID/IL-2Rgcnull (NSG) mice (NCI CCR Animal Resource Program/NCI Biological Testing Branch) were housed under pathogen-free conditions. Mice were inoculated with 4-5 million luciferase-expressing IMR5 or HepG2 cells intravenously or intraperitoneally. After 16-32 days, mice were randomly assigned to groups receiving 5 or 10 million CAR-T cells via intravenous injection.

All mice were regularly monitored for the tumor signal by intraperitoneally injecting 3 mg D-luciferin (PerkinElmer) for 10 minutes, followed by imaging using the Xenogen IVIS Lumina (PerkinElmer). The bioluminescence signal flux was analyzed using the Living Image Software (Revvity) as photons per second per square centimeter per steradian (photons/s/cm^2^/sr).

### Quantification and statistical analysis

Flow cytometry was conducted using the Sony ID7000, and the data were analyzed with FlowJo software. For the mouse model studies, the Xenogen IVIS Lumina (PerkinElmer) was utilized, and the Living Image Software (Revvity) was employed to assess the bioluminescence signal flux. Data analysis was performed using one-way ANOVA with Tukey’s multiple comparisons test or two-way ANOVA with Tukey’s multiple comparisons test, both executed with GraphPad Prism (Dotmatics). Results are presented by the mean and standard deviation (SD), with P values of less than 0.05 considered statistically significant.

## Data availability statement

The data generated in this study are available within the article.

## Acknowledgements

We thank the National Cancer Institute (NCI) Center for Cancer Research (CCR) Animal Resource Program/NCI Biological branch for providing the NSG mice used in the present study, and NCI CCR/Leidos Animal Facility for assisting in animal support. We thank Swati Priya in the NCI and the NIH Fellows Editorial Board for editorial assistance.

## Author contributions

M.Ho, as the guarantor of this study, conceived the research project and supervised the research. M.Ho designed the studies. M.Huo and M.Ho wrote the manuscript. M.Huo, D.L., N.L., A.Q., D.H., J.S., M.P., L.H., K.M., and M.E. performed the experiments. D.H., J.S., M.P., L.H., K.M., and M.E. produced the allogeneic CAR-T cells, and M.Ho provided funding and resources. All the authors reviewed, edited, and approved the manuscript.

